# Nucleotide substitutions during speciation may explain substitution rate variation

**DOI:** 10.1101/2020.08.19.256891

**Authors:** Thijs Janzen, Folmer Bokma, Rampal S. Etienne

**Affiliations:** Groningen Institute for Evolutionary Life Sciences, University of Groningen, Box 11103, 9700 CC Groningen, The Netherlands; Center for Ecological and Evolutionary Synthesis (CEES) Department of BioSciences, University of Oslo, PO Box 1066, Blindern, 0316 Oslo, Norway

**Keywords:** molecular clock, speciation, phylogenetic reconstruction, substitution rate variation

## Abstract

Although molecular mechanisms associated with the generation of mutations are highly conserved across taxa, there is widespread variation in mutation rates between evolutionary lineages. When phylogenies are reconstructed based on nucleotide sequences, such variation is typically accounted for by the assumption of a relaxed molecular clock, which is just a statistical distribution of mutation rates without much underlying biological mechanism. Here, we propose that variation in accumulated mutations may be partly explained by an elevated mutation rate during speciation. Using simulations, we show how shifting mutations from branches to speciation events impacts inference of branching times in phylogenetic reconstruction. Furthermore, the resulting nucleotide alignments are better described by a relaxed than by a strict molecular clock. Thus, elevated mutation rates during speciation potentially explain part of the variation in substitution rates that is observed across the tree of life.

## INTRODUCTION

Phenotypic diversification occurs at a higher rate in some clades than in others (Simpson 1945; van Valen 1985; Ricklefs 2006; Rabosky et al. 2007; Jansson and Davies 2008) and similarly, there is substantial variation across evolutionary lineages in the rate of molecular evolution (King and Wilson 1975), such as that of nucleotide sequences (Nabholz et al. 2008; Bromham 2011; Dowle et al. 2013; Sung et al. 2016). As a consequence, studies attempting to reconstruct the phylogeny of a clade often find that the sequence data do not support the assumption of a strict molecular clock, i.e. constant substitution rates across lineages. For such cases, phylogenetic inference software allows one to use a relaxed molecular clock (Drummond et al. 2006; Lepage et al. 2007), which assumes that the substitution rate varies between lineages according to a statistical distribution such as a gamma or lognormal distribution. However, the relaxed molecular clock thus introduces at least one additional degree of freedom, namely the variance of the distribution of substitution rates (although some argue that an uncorrelated relaxed clock in effect adds one additional degree of freedom *per branch* (Dornburg et al. 2012; Zhang and Drummond 2020; Douglas et al. 2021)). Moreover, the relaxed clock is a rather ad-hoc solution with little underlying biological reasoning (but see Lartillot and Poujol 2014; Lartillot et al. 2016; Saclier et al. 2018).

A first formal test to detect the impact of speciation on sequence evolution was formulated by Avise and Ayala (Avise and Ayala 1975, 1976), who distinguished gradual evolution from “punctuated equilibria” by comparing sequence evolution in species-rich and species-poor clades. Whereas Avise and Ayala found no evidence for increased sequence evolution in species-rich clades, others did, in tetrapods (Mindell et al. 1989, 1990), sauropsids (Eo and DeWoody 2010) and angiosperms (Duchene and Bromham 2013; Bromham et al. 2015). Furthermore, substitution rates have been found to be positively associated with diversification rates (Fontanillas et al. 2007; Eo and DeWoody 2010; Lanfear et al. 2010a, 2010b; Ezard et al. 2013, but see Goldie et al. 2011).

Several biological processes acting at speciation could lead to accelerated sequence evolution, including, but not limited to founder effects, bottlenecks, inbreeding, hybridization, selection for an increased mutation rate, divergent selection and local adaptation (Venditti and Pagel 2010). Here, we explore how such processes driving sequence evolution during speciation events might affect phylogenetic reconstruction; we posit that differences in apparent substitution rates between lineages are due to processes acting exclusively or predominantly during speciation. Due to (effectively) random extinction of lineages, different branches of a reconstructed phylogeny will differ in how often they experienced such short episodes of accelerated substitution rates, resulting in differences in apparent substitutions rates along these branches. Our approach is two-fold: first, we explore whether inclusion of substitutions during speciation affects phylogenetic inference, and, if so, which aspects of the inferred phylogenetic tree are affected. Second, we explore whether substitutions during speciation can explain variation in estimated substitution rates.

## METHODS

We propose a model where substitutions accumulate not only along the branches of a phylogeny, but also at speciation events, including not only the internal nodes of the phylogeny but also those pruned from the phylogeny by extinction. We first make the standard assumption that gradual sequence evolution along a phylogenetic branch can be modeled as a time-homogeneous Markov process with substitution matrix:

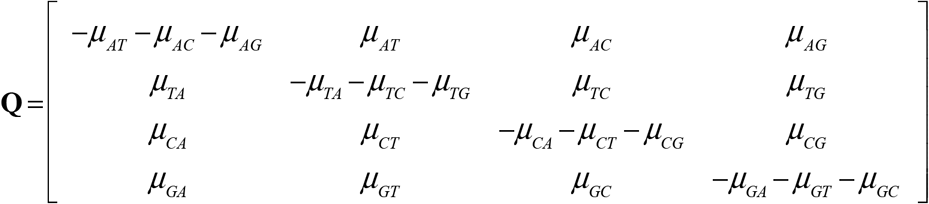

where *μ*_*ij*_ denotes the mutation rate from nucleotide *i* to nucleotide *j*. The transition probabilities of nucleotide substitutions after time *t* of gradual sequence evolution are then given by the matrix

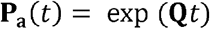

where the subscript **a** indicates anagenetic change, i.e. gradual accumulation of substitutions over time. This matrix can be multiplied with an initial probability vector at time *t* = 0 to yield the probabilities for each of the four nucleotides at time *t*.

In addition to gradual sequence evolution over time, we assume that sequences may change rapidly during speciation. We can thus assume another matrix **P**_**c**_ (subscript c for “cladogenetic”) that describes the nucleotide transition probabilities during a single speciation event.

The processes that may accelerate sequence evolution during speciation, such as founder effects, bottlenecks, inbreeding, hybridization, and adaptation to novel environments, may well result in different kinds of substitutions than those that take place over time in established species. However, for mathematical convenience we will here assume that we can write:

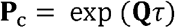

Where τ is a parameter that measures the effect of substitutions during speciation. In other words, we assume that nucleotide sequence evolution is only accelerated during speciation events, but not qualitatively altered: the *μ*_*ij*_ used in **P**_c_ must be identical to those used in **P**_**a**_. The acceleration is then measured by parameter τ: a single speciation event causes as much sequence evolution as τ years of gradual evolution over time within each lineage. Thus, larger values of τ correspond to a larger experienced effect at the nodes, similar to sequence evolution along a branch of length τ. For τ = 0, **P**_c_ becomes the identity matrix, and our model reduces to the standard model of sequence evolution that only assumes substitutions along phylogenetic branches. Important to note here is that both daughter lineages resulting from a speciation event experience substitutions independently (see the Supplement for a model where substitutions in both daughter lineages are dependent on each other). Furthermore, we emphasize that we assume the speciation process to happen in a similar fashion across a tree, assuming an identical τ for all nodes in the tree. Later versions of the model could potentially relax this assumption, provided independent information about speciation dynamics.

For simplicity we assume in our simulations that sequence evolution can be modeled as a Jukes-Cantor process (Jukes and Cantor 1969), for which **Q** is given by:

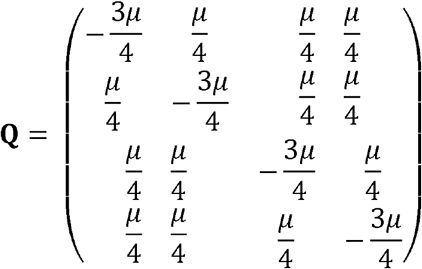

Several existing software packages (e.g. the R package phangorn, Schliep 2011; the python module pyvolve, Sipos *et al*. 2011; the R package phylosim, Spielman & Wilke 2015) provide algorithms to simulate sequence evolution along the branches of the phylogeny, given a rooted phylogeny and a root sequence (e.g. some arbitrary sequence assumed to represent the ancestral sequence), by applying the transition matrix sequentially along the phylogenetic tree. Here, we extend this methodology to also include substitutions accumulated at the nodes of the phylogeny. We implemented this in the R package ‘nodeSub’ (https://github.com/thijsjanzen/nodeSub, and soon available on CRAN).

### Testing the impact of node substitution models using simulations

To identify the amount of error in phylogenetic inference caused by assuming a (relaxed) molecular clock when substitutions actually arise (in part) during speciation, we simulated sequence evolution on known trees and then reconstructed the phylogeny from the simulated sequences, assuming strict and relaxed molecular clocks. We simulated sequence evolution with the node substitution model introduced above, with various degrees of sequence accumulation at the nodes of the tree (*τ*), and with various extinction rates. We then compared the resulting trees with the original true tree using a number of statistics: the gamma statistic (Pybus and Harvey 2000), the beta statistic (Aldous 2001), the mean branch length (Faith 1992; Clarke and Warwick 2001), crown age, the normalized Lineages Through Time (nLTT) statistic (Janzen et al. 2015) and the Jenson-Shannon distance metric comparing the Laplacian spectrum (Lewitus and Morlon 2016).

Phylogenetic reconstruction was performed with BEAST2 (Bouckaert et al. 2019) using the R package *babette* (Bilderbeek and Etienne 2018). BEAST2 inference was performed using default priors, with a birth-death prior as tree prior (or a Yule prior if the extinction rate was zero), the Jukes-Cantor nucleotide substitution model, and a strict or relaxed clock model. The BEAST chain was run for 10 million steps, whilst sampling a tree every 5000 steps. After completion, the first 10% of the chain was discarded as burn-in.

### Assessing error in phylogenetic reconstruction: the twin tree

Errors observed when comparing with the true tree include both errors incurred by the node substitution model chosen, and errors accumulated in the phylogenetic inference process even when the models used in inference are identical to those generating the data (e.g. stochasticity in substitution accumulation, stochasticity in phylogenetic tree creation). Furthermore, additional effects arising during alignment simulation might affect our findings, such as the impact of parameter values (sequence length, substitution rate, birth rate, death rate), and of multiple substitutions at the same site (the node-density-effect) as well as potential biases or interactions between summary statistics. To correct for these effects, so as to isolate the error induced by using a node substitution model from other sources of error, we inferred a phylogenetic tree for a twin alignment (*sensu* Bilderbeek, Laudanno & Etienne 2020). This twin alignment has exactly the same number of substitutions and is based on the same true tree, but instead of using a node substitution model to generate the alignment, it results from using either a strict-clock or relaxed-clock substitution model. Using this twin alignment, we performed phylogenetic reconstruction with BEAST2 as for the original alignment, and estimated the same summary statistics for the posterior distribution of trees. The error introduced by the node substitution model is then the difference between the error of the node substitution posterior and the error in the twin posterior. In summary, we use this twin approach as a control treatment, in order to correct for all potential sources of additional error other than that of our proposed substitution model.

### Obtaining a twin alignment

We generated a twin alignment conditional on a phylogeny, a node substitution model, and a mutation rate. Because an alignment generated using a node substitution model (with *τ* > 0) has accumulated substitutions at the nodes in addition to those along the branches, the overall number of substitutions accumulated is higher than for an alignment simulated using the same mutation rate and a model with substitutions only on the branches. Thus, in order to generate a *twin* alignment that contains the same amount of information (substitutions) we increased the mutation rate. We did this by calculating the estimated time spent at the nodes, relative to the time spent on the branches, and using this as an estimate of the expected fraction of the number of substitutions on the nodes, relative to the number of substitutions on the branches, assuming that substitutions accumulate at the same rate on both branches and nodes. That is, the mutation rate used in generating the twin alignment is calculated as:

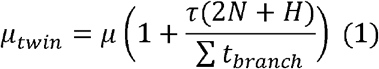

where *μ* is the mutation rate used in the node substitution model, *τ* is the time spent on the node, *N* is the number of internal nodes in the tree, *H* is the number of hidden nodes in the tree and Σ*t*_*branch*_ is the total branch length of the tree. The factor *2N* arises from the independent accumulation of substitutions during a node substitution event for both daughter lineages.

During simulation of node substitution alignments, we kept track of the substitutions accumulated at each node and branch, which allowed us to directly measure the contribution of substitutions accumulated at the nodes (i.e. τ(2*N*+*H*)) relative to those accumulated at the branches (i.e.. Σ*t*_*branch*_) This provided us with an estimate of *μ*_*twin*_, and with an estimate of the total number of substitutions arising during simulation of the alignment. We then used the obtained estimate for *μ*_*twin*_ to generate *<u>twin</u>* alignments, again tracking all substitutions, until we obtained an alignment exactly matching the number of accumulated substitutions of the alignment simulated with the node substitution model.

### Node-density-effect

The method we used to simulate substitutions along branches (and nodes) ignores repeated mutations at the same site, which may lead to a node-density-effect. Because the node-density-effect can mask the effect of node substitutions, we made sure in two distinct ways that our results are not affected by this effect. Firstly, by using a *twin* alignment, any resulting node-density-effects are mirrored in the *twin* alignment as well, ensuring that any additional errors picked up do not reflect errors induced by the node-density-effect. Secondly, we repeated our analysis using a different simulation method that explicitly tracks repeated mutations for the Jukes-Cantor model (see Supplementary Information for details and results). We find that this more explicit simulation method yielded virtually identical results to the more general approach described in the main text.

### Simulation settings

We generated birth-death trees with varying degrees of extinction rate *d* in [0, 0.1, 0.3, 0.5] and a single speciation rate of *b* = 1. Trees were simulated conditional on 100 tips, using the function `sim.bd.taxa` from the R package TreeSim (Stadler 2011). Across all settings, we simulated sequences of 10 kb, with *μ* = 0.001.

### Varying the time spent on the nodes relative to the crown age

We varied *τ* in [0, 0.01, 0.05, 0.1, 0.2, 0.4] times the crown age (e.g. when the crown age of the simulated tree is 3MY, τ = [0, 0.03, 0.15, 0.3, 0.6, 1.2] MY). Again, for each combination of τ and extinction (*d* = [0, 0.1, 0.3, 0.5]) we simulated 100 trees and for each tree we generated one node substitution alignment and one *twin* alignment.

### The impact of tree balance

In unbalanced trees, some terminal branches are connected by many more past branching events to the root of the tree than are other terminal branches. Hence, we expect that balance of a tree might have a substantial effect on the error in phylogenetic inference: less balanced trees are expected to have higher error. To test this, we compared fully balanced (፧= 10.0) with extremely unbalanced “caterpillar” trees (፧= -2). We did so by simulating the branching times of a birth-death tree, and assigning these to a fully balanced or fully unbalanced topology. Thus, the only difference between the trees is the topology. Then, for both the balanced and unbalanced tree a node substitution alignment was generated, with the same number of total substitutions, and setting *τ* as a function of crown age.

As an extra check, we also generated a node substitution alignment for the original birth-death tree from which the branching times were used. For all three alignments we inferred a phylogenetic tree as in the other scenarios, and compared the error in phylogenetic inference. For caterpillar trees with extreme unbalance we were unable to calculate the Laplacian Spectrum, hence we omitted the Laplacian Spectrum summary statistic in this analysis.

### Support for strict and relaxed clock models

To test whether the alignment originating from a process with node substitutions was better described by a relaxed than by a fixed clock model, we repeated the analysis, but now inferred the marginal likelihood of the relaxed clock and strict clock models using the “NS” package for BEAST2, which applies Nested Sampling to obtain the marginal posterior likelihood for both models (Russel et al. 2019). We used the function ‘`bbt_run`’ from the *babette* package (Bilderbeek and Etienne 2018) in combination with the function ‘`create_ns_mcmc`’ from the *beautier* package (Bilderbeek and Etienne 2018). This performs a Nested Sampling MCMC run using BEAST2, which runs until convergence is detected. Then, we converted the obtained marginal likelihoods to a relative weight for each model (by dividing both marginal likelihoods by their sum) which allows for comparison of posterior support for each model across parameter settings and trees.

## RESULTS

### Summary statistics

We compared summary statistics of trees inferred from alignments using the node substitution model, with summary statistics of twin trees inferred from alignments with identical information content, but generated without the node substitution model (e.g. with only substitutions along the branches, and a fixed clock rate). We find that summary statistics that are influenced by branching times are affected (Figure 1, gamma, nLTT statistic, mean branch length and crown age). For these summary statistics, we find an increased difference as the time spent on the nodes *τ* increases. The differences in the values of these summary statistics manifest themselves across all extinction rates, although the effect tends to be somewhat smaller for higher extinction rates.

**Figure 1.**
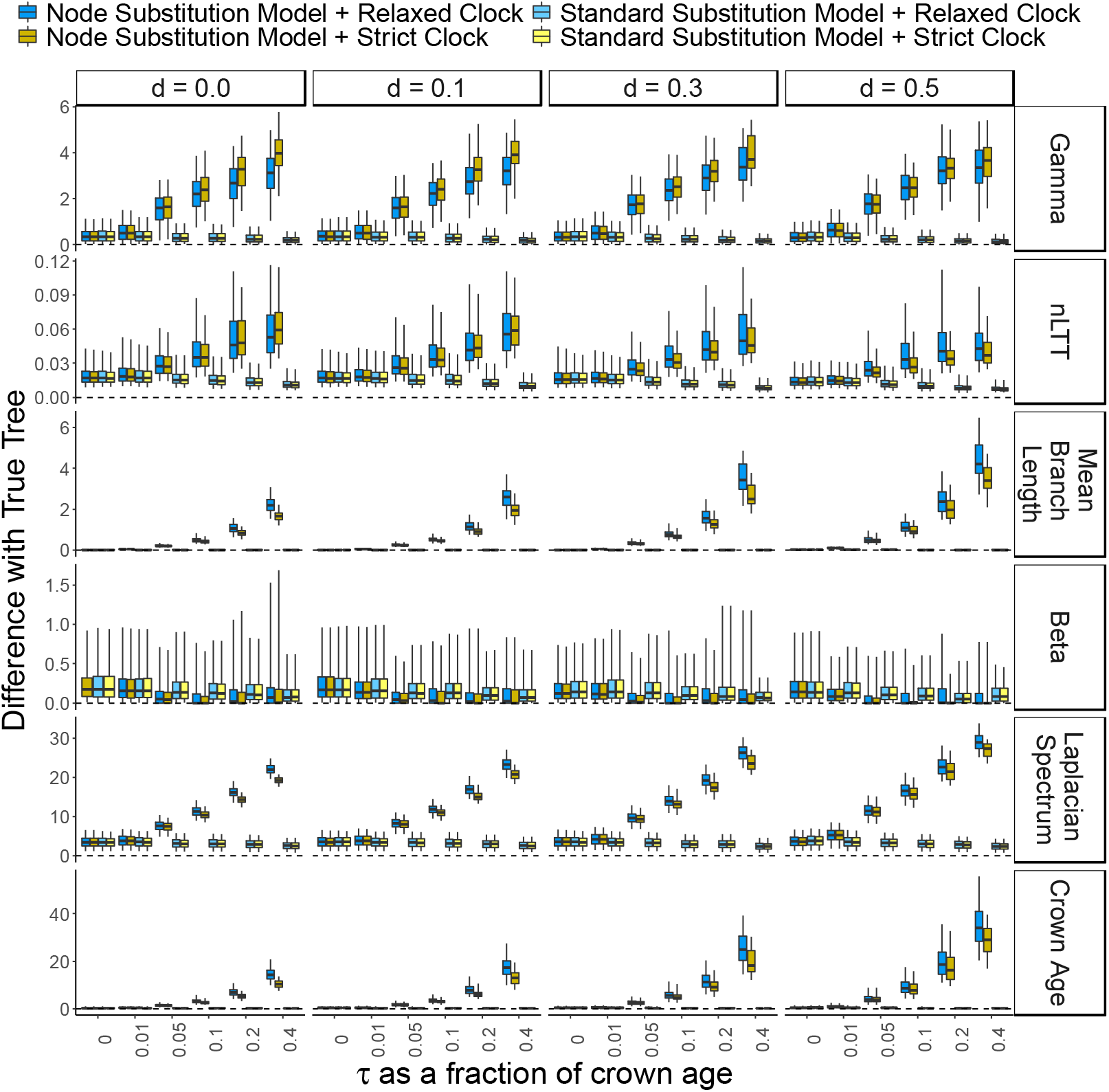
Difference in summary statistic values for trees inferred from an alignment generated with node substitutions, and *twin* trees that were inferred from an alignment generated without node substitutions (grey boxplots), both compared with the summary statistics of the true tree. We explored *τ* (the amount of time spent on each node) as a fraction of crown age (horizontal axis), and the impact of extinction (*d*, columns). The summary statistics are the beta and gamma statistic, Laplacian spectrum, mean branch length, nLTT statistic, and crown age. The figure shows that with increasing *τ*, trees inferred from an alignment generated with node substitutions show larger differences with the true tree than trees inferred from an alignment generated without node substititions. Differences with the true tree are larger for trees inferred using the strict clock model than for those using the relaxed clock model, but only for the alignment generated with node substitutions.

### The impact of tree balance

Tree balance clearly influences the sensitivity of inference to node substitutions (Figure 2). The inference error is larger for unbalanced trees, again only for the gamma and the nLTT statistic. Fully balanced trees show slightly less error than birth-death trees. Overall, all three types of trees show an increased error when alignments are generated using the node substitution model. Errors are particularly large for the beta statistic, but that is expected because it measures topological features of the tree that we modified artificially.

**Figure 2.**
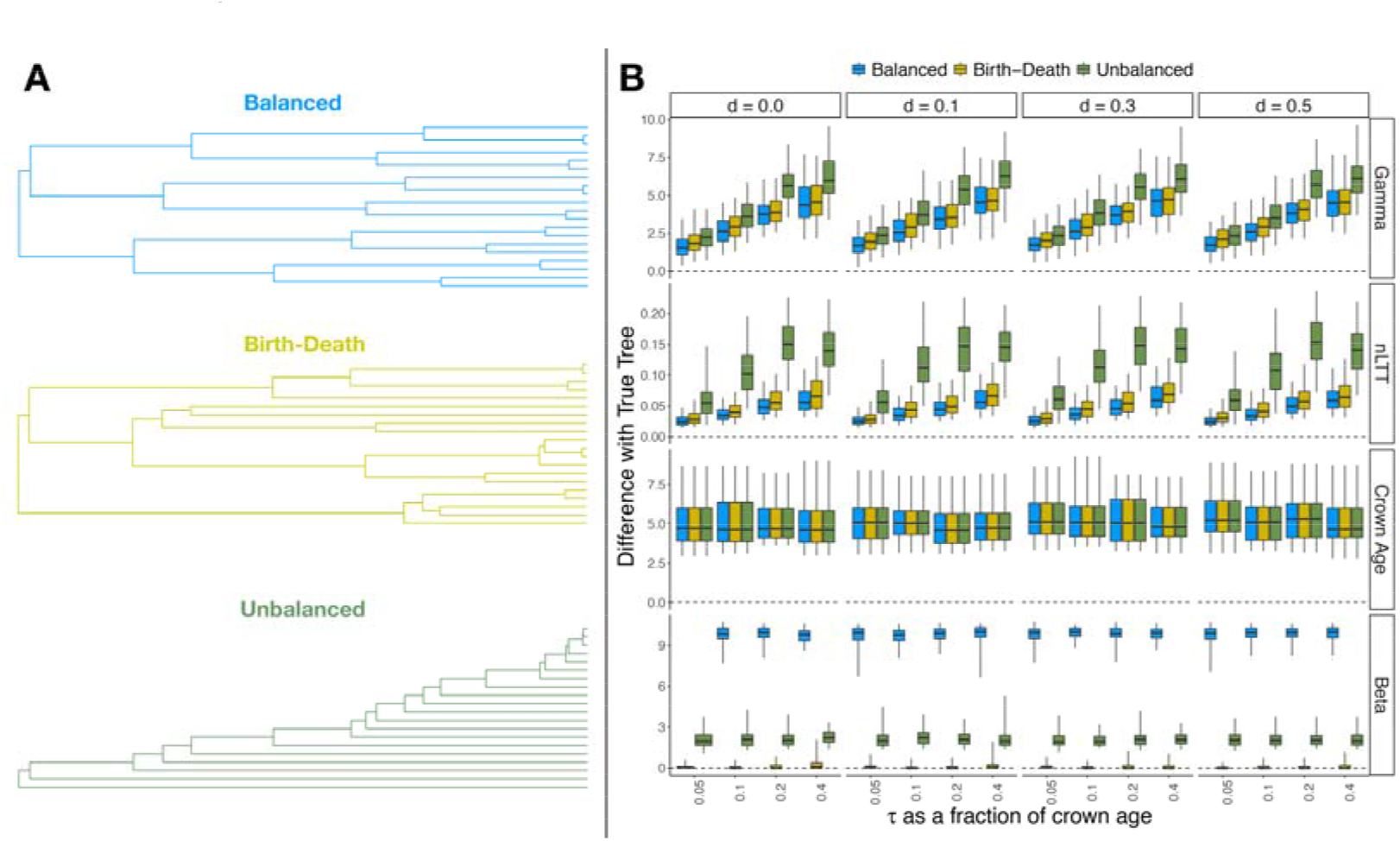
Effect of the node substitution model for phylogenies differing in balance **A)** example plots of a randomly generated birth-death tree (top), a fully balanced tree generated using the same branching times as the birth-death tree (middle) and a very unbalanced tree generated using the same branching times as the birth-death tree (bottom). Shown are trees with 20 tips for illustrative purposes, but results in **B** are from trees with 100 tips. **B)** Difference in summary statistic with the true birth-death tree for phylogenetic trees inferred from alignments generated using the node substitution model on either balanced, unbalanced or random trees. We explore *τ* as a fraction of crown age (horizontal axis), and the impact of extinction (*d, columns*). The dotted line indicates zero difference with the true tree. The summary statistics are the beta and gamma statistic, nLTT statistic and tree height. Balanced and birth-death trees tend to have similar inferred error, whereas unbalanced trees differ strongly, with a much larger error for the gamma and nLTT statistic.

### Support for strict and relaxed clock models

We compared the relative support for each model, reflected by the relative weight of the marginal likelihood. With an increasing amount of time spent at the nodes *τ*, the median weight of the relaxed clock model increases for the node substitution alignment, with generally (across extinction rates) a higher weight than the strict clock model for values of *τ* that are equal or larger than 0.1 times the crown age (Figure 3). For the twin alignment, the strict clock model is preferred, as expected, because this is the generating model.

**Figure 3.**
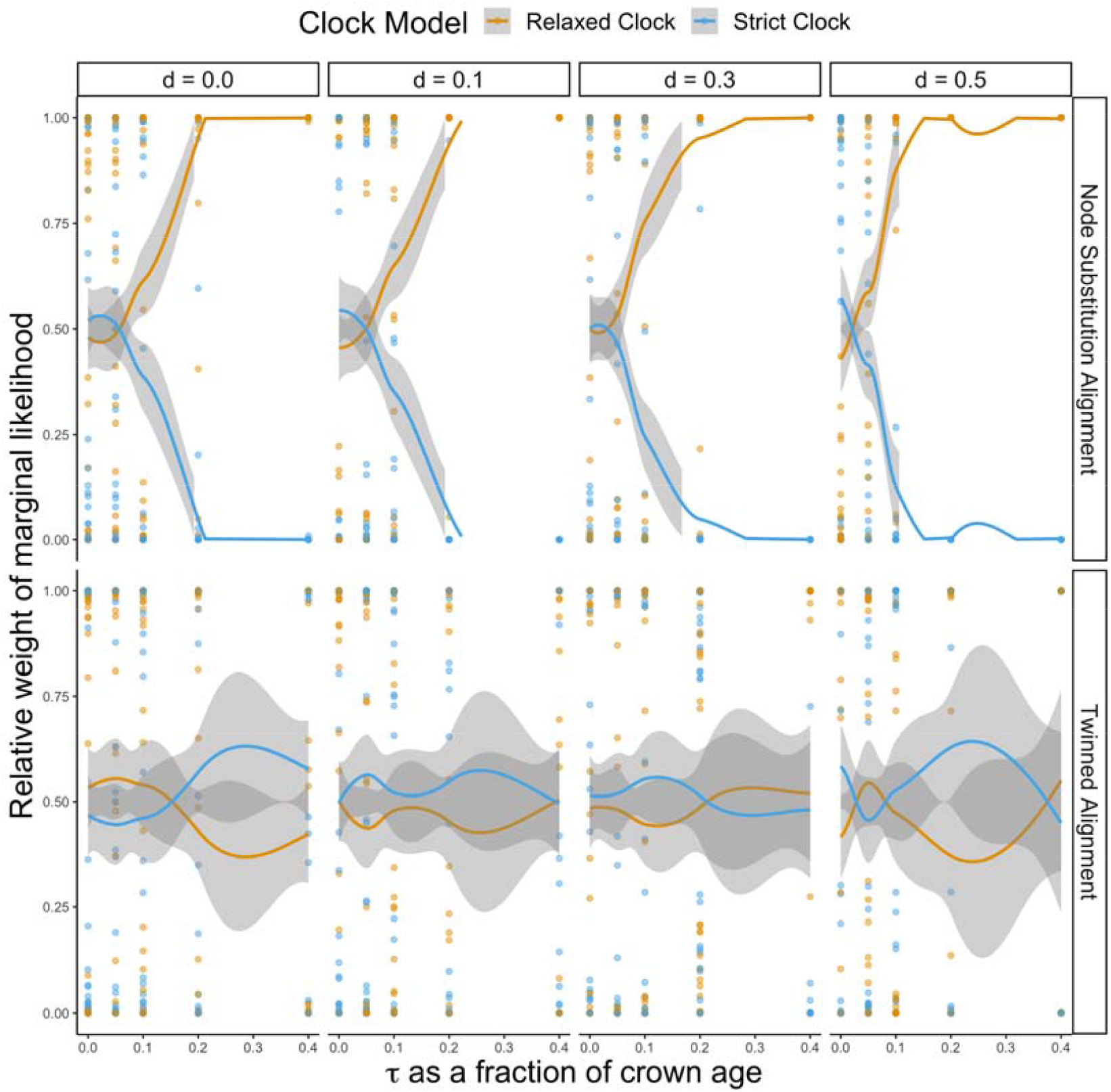
Marginal likelihood weight of the relaxed and strict clock model for varying time spent on the nodes (*τ*), where *τ* ፧is chosen as fraction of the crown age. Alignments generated with a node substitution model (top row) are compared with alignments generated without node substitutions (bottom row). Per parameter combination, 100 replicate trees were analyzed. Because many dots are plotted on top of each other, we use solid lines to indicate the best fitting locally estimated scatterplot smoothing (LOESS), and the 95% Confidence interval (grey shaded area) of the LOESS curve. As the time spent on the nodes increases, posterior support for the relaxed clock model increases, but only if the alignment was generated with a node substitution model.

For low values of *τ* (smaller than 0.1 times the crown age), we do not find any effect of the balance of the tree on the marginal likelihood of the relaxed clock model (Figure 4), in line with our finding above. However, for intermediate values of *τ* (0.1 and 0.2), we find that unbalanced trees tend to have a higher marginal weight for the relaxed clock model. For high values of *τ* (0.4), we find that the marginal weight for the relaxed clock model is always higher, regardless of the balance of the tree.

**Figure 4.**
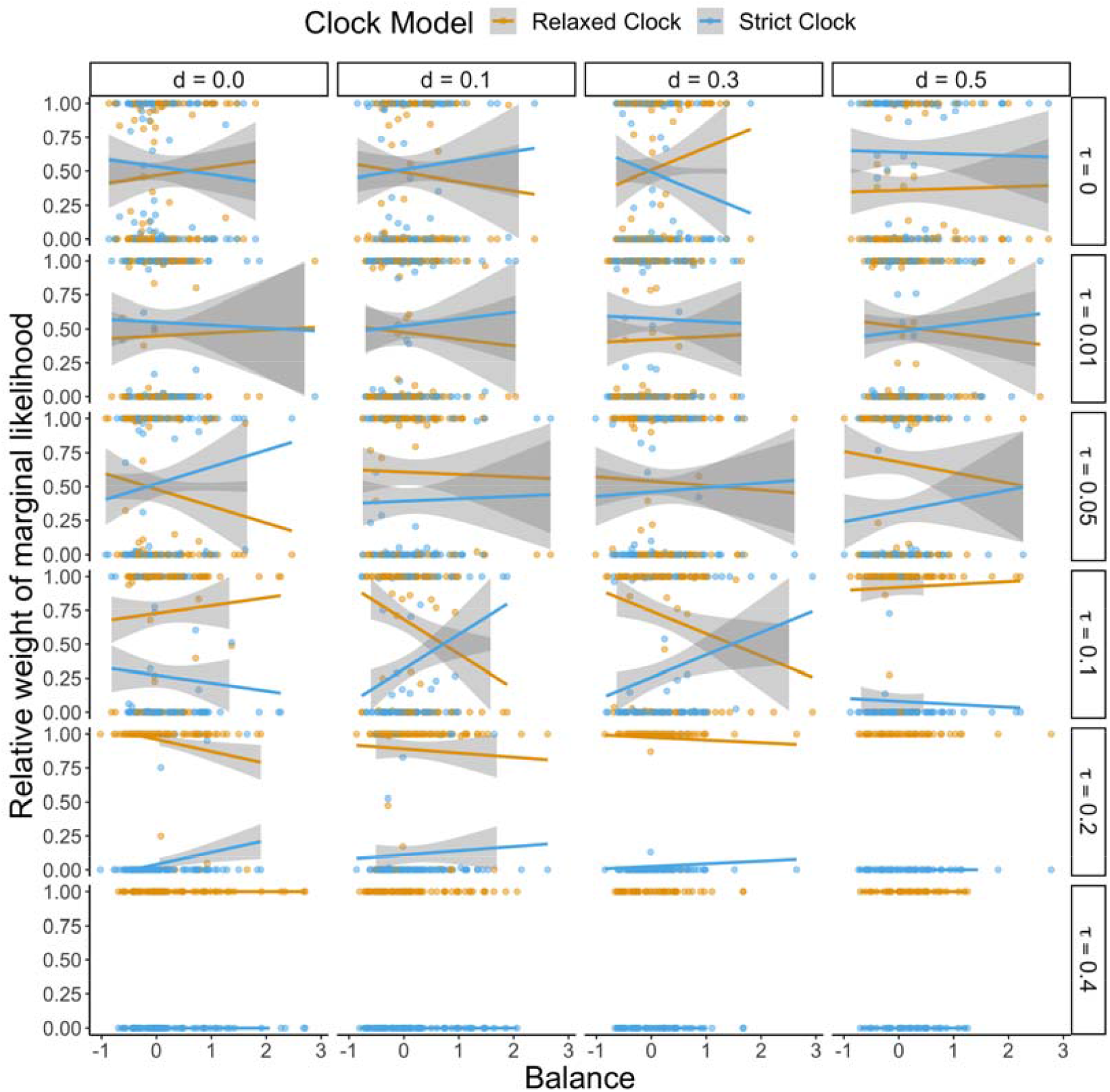
Marginal likelihood weight of the relaxed clock model for trees of varying balance, split out across different extinction rates (d = [0, 0.1, 0.3, 0.5]) and time spent on the nodes (*τ*), where *τ* ፧is chosen as fraction of the crown age (e.g. *τ* = 0.1 reflects a node time of 10% of the crown age). Per parameter combination, Solid lines indicate the best fitting linear regression and the 95% confidence interval (grey shaded area) of regression. With increasing values of *τ*, the relative weight of the relaxed clock model becomes larger. For smaller values of *τ*, the relative weight of the relaxed clock model is negatively correlated with the balance of the tree, with unbalanced trees having a higher relative weight.

## DISCUSSION

We have shown that an increased substitution rate during speciation events potentially provides a mechanistic explanation of variation in substitution rates across the branches of phylogenetic trees. Trees inferred from alignments generated with this substitution model differ substantially from trees inferred from alignments generated with a standard substitution model, especially concerning branching times. Furthermore, we find that this new substitution model can potentially explain widespread support for relaxed molecular clocks.

If sequence evolution mainly occurs during speciation, this would lead to a correlation between species richness and substitution rate. However, this correlation could also be an artifact of phylogenetic reconstruction known as the node-density-effect (Fitch and Bruschi 1987; Fitch and Beintema 1990). The node-density-effect reflects the inability to detect multiple mutations occurring at the same site, thus causing an underestimate of the true branch length, especially for longer branches where the probability of multiple mutations occurring at the same site is higher. Because species-rich parts of phylogenies tend to have shorter branches, sequence evolution in these species-rich parts is less underestimated than in species-poor parts, causing a correlation between the number of observed substitutions and species diversity. Pagel et al. tested for the impact of speciation events, and of the node-density-effect in 122 phylogenies, spanning 4 taxa (Pagel et al. 2006). Using previously demonstrated methodology to detect the node-density effect (Webster et al. 2003; Venditti et al. 2006), they showed that in 57 of the examined phylogenies, they could detect a signature of increased sequence evolution during speciation events. However, this was the result of the node-density effect in 22 out of these 57 phylogenies. Here, disentangling sequence evolution during speciation from confounding factors such as the node-density effect, but also stochasticity in tree simulation, stochasticity during alignment simulation and error or bias in tree inference, has proven to be a non-trivial endeavor. In order to assess the impact of node substitutions, we therefore separated error due to assuming an alternative substitution model from the errors introduced by the factors mentioned above. To do so we extended the twinning approach (introduced by Bilderbeek, Laudanno and Etienne (2020)) to assess the impact of choosing a different tree prior) to explore the impact of a different substitution model. The twinning approach succeeds by replicating the chosen analysis pipeline, but using *control* data that have been generated using known models and priors. The impact of the node substitution model then follows from the difference between results obtained with the node substitution model and results obtained with the *twin* (control) pipeline: errors are then due to model misspecification, and not stochastic uncertainty produced by the analysis pipeline.

One might expect that a high extinction rate, by elevating numbers of hidden nodes, would lead to a greater impact of node substitutions. It may therefore be counterintuitive that in our simulation study we did not find such an effect of higher rates of extinction. However, we conditioned our alignments on the same total number of substitutions, to ensure that alignments with and without node substitutions contained the same information content. Thus, with higher extinction and hence more hidden nodes, fewer substitutions occur on the observed nodes. Because the number of hidden nodes is proportional to branch length (Eq.2), it is interpreted as substitutions on the branches. Consequently, higher extinction rates erase the signature of node substitutions. Potentially, this provides a way to distinguish between phylogenetic models: although every constant-rate birth-death model has a corresponding zero-extinction model with a time-varying speciation rate that yields the same probability of the reconstructed tree (Nee et al. 1994; Louca and Pennell 2020), the resulting alignments under the node substitution model will not be similar. Because the birth-death tree includes extinction events, substitution patterns will be different from those of the tree generated with the time-varying speciation rate model.

Distinguishing phylogenetic models will become more feasible if some of the simplifying assumptions made here are relaxed. The model we propose here takes the simplest form, assuming a Jukes-Cantor (Jukes and Cantor 1969) substitution matrix, identical substitution rates, identical substitution matrices between nodes and branches, and constant birth-death rates over time. These assumptions were made as a most basic starting point, but can be relaxed in future analyses, for instance by introducing a different substitution matrix at the nodes, or by studying the effect of node substitutions on trees that are generated by diversity-dependent speciation rates (Etienne et al. 2012). By starting with the most tractable version of the node substitution model, we have provided a first proof of concept of the potential impact of node substitutions without overcomplicating matters.

Previous methods have applied rather ad-hoc corrections to account for differences in substitution rates across different branches in the same phylogeny, typically referred to as the ‘relaxed clock’ approach. These methods provide satisfying statistical solutions to account for variation in substitution rates, but refrain from providing biological explanations for this observed phenomenon. The node substitution model we introduce here provides this explanation: branches that have accumulated a number of ‘hidden’ branching events, e.g. speciation events of species that have subsequently gone extinct, have a higher number of accumulated substitution events during these ‘hidden’ speciation events. When we compared the marginal likelihood of the relaxed clock model versus the strict clock model for alignments generated with the node substitution model, we found that marginal likelihoods for the relaxed clock model are much higher. This indicates that our proposed process of accumulating substitutions during speciation events can generate patterns in the alignment that are picked up by phylogenetic methods as evidence for a relaxed clock model, without actually using a relaxed clock model.

The notion of accelerated evolution during speciation events ties in closely with the theory of punctuated equilibrium; where Eldredge and Gould (1972) proposed that evolution perhaps is not a gradual process, but rather a process with distinct bursts of phenotypic and morphological change. Their theory was influenced by ideas like Lerner’s “genetic homeostasis” (Lerner 1954), which had earlier inspired Mayr (1954) to suggest that the formation of new species involves “genetic revolutions”. Our framework provides a step towards being able to test this notion, where information on the estimated fraction of substitutions accumulated at the nodes can directly inform us about whether the majority of substitutions is accumulated over long periods of time in established lineages (e.g. along branches), or during speciation (e.g. at the nodes).

The present study aims to demonstrate that substitutions accumulated during speciation might explain the prevalence of the relaxed molecular clock in phylogenetic analysis. We found that substitutions during speciation may profoundly affect phylogenetic inference: if node substitutions are not taken into account, branching times tend to be overestimated, even when a relaxed clock is used to counteract the effect of “hidden nodes”. This suggests that incorporation of a node substitution model improves phylogenetic inference. However, the computation of the likelihood of our model (and estimation of associated τ values), is non-trivial, and might escape computational feasibility due to the high state space required. Manceau *et al*. (2020) have taken a first step towards formulating such a likelihood. They introduced an alternative solution for punctuated equilibrium-like patterns in molecular evolution through the implementation of spikes of substitution, e.g. moments in time at which there is an increased substitution rate. They let these moments occur at speciation events, and also model the probability of such an event happening at a speciation event (rather than assuming that they always occur, like we did here). However, they have to assume both the topology and branching times to be fixed. This provides a promising first step, but given that the number of hidden nodes depends on the diversification rates, what is ultimately needed is a new method that jointly infers the diversification model and the substitution model.

With our introduction of the node substitution model, we hope to stimulate discussion on the biological explanation of variation in substitution rates within and across phylogenies. Furthermore, we hope to have set a first step in improving our understanding of this variation, and improving phylogenetic inference as a whole.

## ACKNOWLEDGEMENTS

We thank Sebastian Hoehna and three anonymous referees for providing comments on the manuscript. FB was supported by the Research Council of Norway, grant 263149. RSE thanks the Netherlands Organization for Scientific Research (NWO) for financial support through VIDI and VICI grants. TJ thanks the Center for Information Technology of the University of Groningen for their support and for providing access to the Peregrine high performance computing cluster.

## References

Aldous D. 2001. Stochastic models and descriptive statistics for phylogenetic trees, from Yule to today. Stat. Sci. 16:23–34.

Avise J.C., Ayala F.J. 1975. Genetic change and rates of cladogenesis. Genetics. 81:757–773.

Avise J.C., Ayala F.J. 1976. Genetic Differentiation in Speciose Versus Depauperate Phylads: Evidence from the California Minnows. Evolution. 30:46.

Bilderbeek R.J.C., Etienne R.S. 2018. babettel፧: BEAUti 2, BEAST2 and Tracer for R. 2018:2034–2040.

Bilderbeek R.J.C., Laudanno G., Etienne R.S. 2020. Quantifying the impact of an inference model in Bayesian phylogenetics. Methods Ecol. Evol.:1–8.

Bouckaert R., Vaughan T.G., Barido-Sottani J., Duchene S., Fourment M., Gavryuskina A., Heled J., Jones G., Kuhnert D., De Maio N., Matschiner M., K. Mendes F., Muller N.F., Ogilvie H.A., du Plessis L., Popinga A., Rambaut A., Rasmussen D., Siveroni I., Suchard M.A., Wu C.H., Xie D., Zhang C., Stadler T., Drummond A.J. 2019. BEAST 2 .5፧: An advanced software platform for Bayesian evolutionary analysis. PLoS Comput. Biol. 15:e1006650.

Bromham L. 2011. The genome as a life-history character: Why rate of molecular evolution varies between mammal species. Philos. Trans. R. Soc. B Biol. Sci. 366:2503–2513.

Bromham L., Hua X., Lanfear R., Cowman P.F. 2015. Exploring the relationships between mutation rates, life history, genome size, environment, and species richness in flowering plants. Am. Nat. 185:508–524.

Clarke K., Warwick R. 2001. A further biodiversity index applicable to species lists: variation in taxonomic distinctness. Mar. Ecol. Prog. Ser. 216:265–278.

Dornburg A., Brandley M.C., McGowen M.R., Near T.J. 2012. Relaxed clocks and inferences of heterogeneous patterns of nucleotide substitution and divergence time estimates across whales and dolphins (Mammalia: Cetacea). Mol. Biol. Evol. 29:721–736.

Douglas J., Zhang R., Bouckaert R. 2021. Adaptive dating and fast proposals: Revisiting the phylogenetic relaxed clock model. PLOS Comput. Biol. 17:e1008322.

Dowle E.J., Morgan-Richards M., Trewick S.A. 2013. Molecular evolution and the latitudinal biodiversity gradient. Heredity (Edinb). 110:501–510.

Drummond A.J., Ho S.Y.W., Phillips M.J., Rambaut A. 2006. Relaxed phylogenetics and dating with confidence. PLoS Biol. 4:e88.

Duchene D., Bromham L. 2013. Rates of molecular evolution and diversification in plants: Chloroplast substitution rates correlate with species-richness in the Proteaceae. BMC Evol. Biol. 13:1.

Eldredge N., Gould S.J. 1972. Punctuated equilibria: an alternative to phyletic gradualism. Models In Paleobiology. Freeman Cooper and Co. p. 82–115.

Eo S.H., DeWoody J.A. 2010. Evolutionary rates of mitochondrial genomes correspond to diversification rates and to contemporary species richness in birds and reptiles. Proc. R. Soc. B Biol. Sci. 277:3587–3592.

Etienne R.S., Haegeman B., Stadler T., Aze T., Pearson P.N., Purvis A., Phillimore A.B. 2012. Diversity-dependence brings molecular phylogenies closer to agreement with the fossil record. Proc. R. Soc. B Biol. Sci. 279:1300–1309.

Ezard T.H.G., Thomas G.H., Purvis A. 2013. Inclusion of a near-complete fossil record reveals speciation-related molecular evolution. Methods Ecol. Evol. 4:745–753.

Faith D.P. 1992. Conservation evaluation and phylogenetic diversity. Biol. Conserv. 61:1–10.

Fitch W.M., Beintema J.J. 1990. Correcting parsimonious trees for unseen nucleotide substitutions: The effect of dense branching as exemplified by ribonuclease. Mol. Biol. Evol. 7:438–443.

Fitch W.M., Bruschi M. 1987. The evolution of prokaryotic ferredoxins--with a general method correcting for unobserved substitutions in less branched lineages. Mol. Biol. Evol. 4:381–394.

Fontanillas E., Welch J.J., Thomas J.A., Bromham L. 2007. The influence of body size and net diversification rate on molecular evolution during the radiation of animal phyla. BMC Evol. Biol. 7:1–12.

Goldie X., Lanfear R., Bromham L. 2011. Diversification and the rate of molecular evolution: No evidence of a link in mammals. BMC Evol. Biol. 11:286.

Jansson R., Davies T.J. 2008. Global variation in diversification rates of flowering plants: Energy vs. climate change. Ecol. Lett. 11:173–183.

Janzen T., Höhna S., Etienne R.S.R.S. 2015. Approximate Bayesian Computation of diversification rates from molecular phylogenies: introducing a new efficient summary statistic, the nLTT. Methods Ecol. Evol. 6:566–575.

Jukes T., Cantor C. 1969. Evolution of protein molecules. Mamm. Protein Metab. 21.

King M.C., Wilson A.C. 1975. Evolution at two levels in humans and chimpanzees. Science. 188:107–116.

Lanfear R., Ho S.Y.W., Love D., Bromham L. 2010a. Mutation rate is linked to diversification in birds. Proc. Natl. Acad. Sci. U. S. A. 107:20423–20428.

Lanfear R., Welch J.J., Bromham L. 2010b. Watching the clock: Studying variation in rates of molecular evolution between species. Trends Ecol. Evol. 25:495– 503.

Lartillot N., Phillips M.J., Ronquist F. 2016. A mixed relaxed clock model. Philos. Trans. R. Soc. B Biol. Sci. 371.

Lartillot N., Poujol R. 2014. Correlated evolution of substitution rates and quantitative traits..

Lepage T., Bryant D., Philippe H., Lartillot N. 2007. A general comparison of relaxed molecular clock models. Mol. Biol. Evol. 24:2669–80.

Lerner I.M. 1954. Genetic Homeostasis..

Lewitus E., Morlon H. 2016. Characterizing and comparing phylogenies from their laplacian spectrum. Syst. Biol. 65:495–507.

Louca S., Pennell M.W. 2020. Extant timetrees are consistent with a myriad of diversification histories. Nature. 580:502–505.

Manceau M., Marin J., Morlon H., Lambert A. 2020. Model-based inference of punctuated molecular evolution. Mol. Biol. Evol.:msaa 144.

Mayr E. 1954. Change of genetic environment and evolution. Evol. as a Process.:157–180.

Mindell D.P., Sites J.W., Graur D. 1989. Speciational Evolution: a Phylogenetic Test With Allozymes in Sceloporus (Reptilia). Cladistics. 5:49–61.

Mindell D.P., Sites J.W., Graur D. 1990. Mode of allozyme evolution: Increased genetic distance associated with speciation events. J. Evol. Biol. 3:125–131.

Nabholz B., Glémin S., Galtier N. 2008. Strong variations of mitochondrial mutation rate across mammals - The longevity hypothesis. Mol. Biol. Evol. 25:120–130.

Nee S., May R.M., Harvey P.H., Trans P., Lond R.S. 1994. The reconstructed evolutionary process. Philos. Trans. R. Soc. Lond. B. Biol. Sci. 344:305–11.

Pagel M., Venditti C., Meade A. 2006. Large punctuational contribution of speciation to evolutionary divergence at the molecular level. Science. 314:119–121.

Pybus O., Harvey P. 2000. Testing macro–evolutionary models using incomplete molecular phylogenies. Proc. R. Soc. B Biol. Sci. 267:2267–72.

Rabosky D.L., Donnellan S.C., Talaba A.L., Lovette I.J. 2007. Exceptional among-lineage variation in diversification rates during the radiation of Australia’s most diverse vertebrate clade. Proc. R. Soc. B Biol. Sci. 274:2915–2923.

Ricklefs R.E. 2006. Global variation in the diversification rate of passerine birds. Ecology. 87:2468–2478.

Russel P.M., Brewer B.J., Klaere S., Bouckaert R.R. 2019. Model Selection and Parameter Inference in Phylogenetics Using Nested Sampling. Syst. Biol. 68:219–233.

Saclier N., François C.M., Konecny-Dupre L., Lartillot N., Guéguen L., Duret L., Malard F., Douady C.J., Lefébure T. 2018. Life history traits impact the nuclear rate of substitution but not the mitochondrial rate in isopods. Mol. Biol. Evol. 35:2900–2912.

Schliep K.P. 2011. phangornl፧: phylogenetic analysis in R. 27:592–593.

Simpson G.G. 1945. SECTION OF BIOLOGY: Tempo and Mode in Evolution. Trans. N. Y. Acad. Sci. 8:45–60.

Sipos B., Massingham T., Jordan G.E., Goldman N. 2011. PhyloSim - Monte Carlo simulation of sequence evolution in the R statistical computing environment..

Spielman S.J., Wilke C.O. 2015. Pyvolvel⍰: A Flexible Python Module for Simulating Sequences along Phylogenies. :1–7.

Stadler T. 2011. Simulating trees with a fixed number of extant species. Syst. Biol. 60:676–684.

Sung W., Ackerman M.S., Dillon M.M., Platt T.G., Fuqua C., Cooper V.S., Lynch M. 2016. Evolution of the insertion-deletion mutation rate across the tree of life. G3 Genes, Genomes, Genet. 6:2583–2591.

van Valen L.M. 1985. Why and how do mammals evolve unusually rapidly. Evol. Theory. 7:127–132.

Venditti C., Meade A., Pagel M. 2006. Detecting the node-density artifact in phylogeny reconstruction. Syst. Biol. 55:637–643.

Venditti C., Pagel M. 2010. Speciation as an active force in promoting genetic evolution. Trends Ecol. Evol. 25:14–20.

Webster A.J., Payne R.J.H., Pagel M. 2003. Molecular phylogenies link rates of evolution and speciation. Science. 301:478.

Zhang R., Drummond A. 2020. Improving the performance of Bayesian phylogenetic inference under relaxed clock models. BMC Evol. Biol. 20:1–28.

