## Supplementary Material for "Nucleotide substitutions during speciation may explain substitution rate variation"

Supplementary information

*Varying the fraction of substitutions on the nodes*

The fraction $f_c$ is given by:

$$f_c = \frac{\tau(2N + H)}{\tau(2N + H) + \sum_i b_i} \quad (S1)$$

where $N$ indicates the number of internal nodes of the tree, $H$ indicates the number of hidden nodes, $b_i$ indicates the length of branch $i$ and $\tau$ indicates the time spent on the node.

Firstly, we verified correctness of equation S1 by keeping track of accumulated branches during simulating alignments. We then a posteriori estimated the observed fraction of substitutions accumulated at the nodes, for a sequence of 10000 bp, with a substitution rate of 1e-3, for d in [0, 0.1, 0.3, 0.5], $\tau$ in [0, 0.01, 0.05, 0.1, 0.2, 0.4] times the crown age and using 100 replicates per parameter combination. Results in figure S1 show that the expected fraction of substitution on the nodes closely follows the observed fraction. Furthermore, the used values of $\tau$ (which align with the values of $\tau$ used in the main text) explore a range of $f_c$ in [0, 0.75], which seems reasonable.

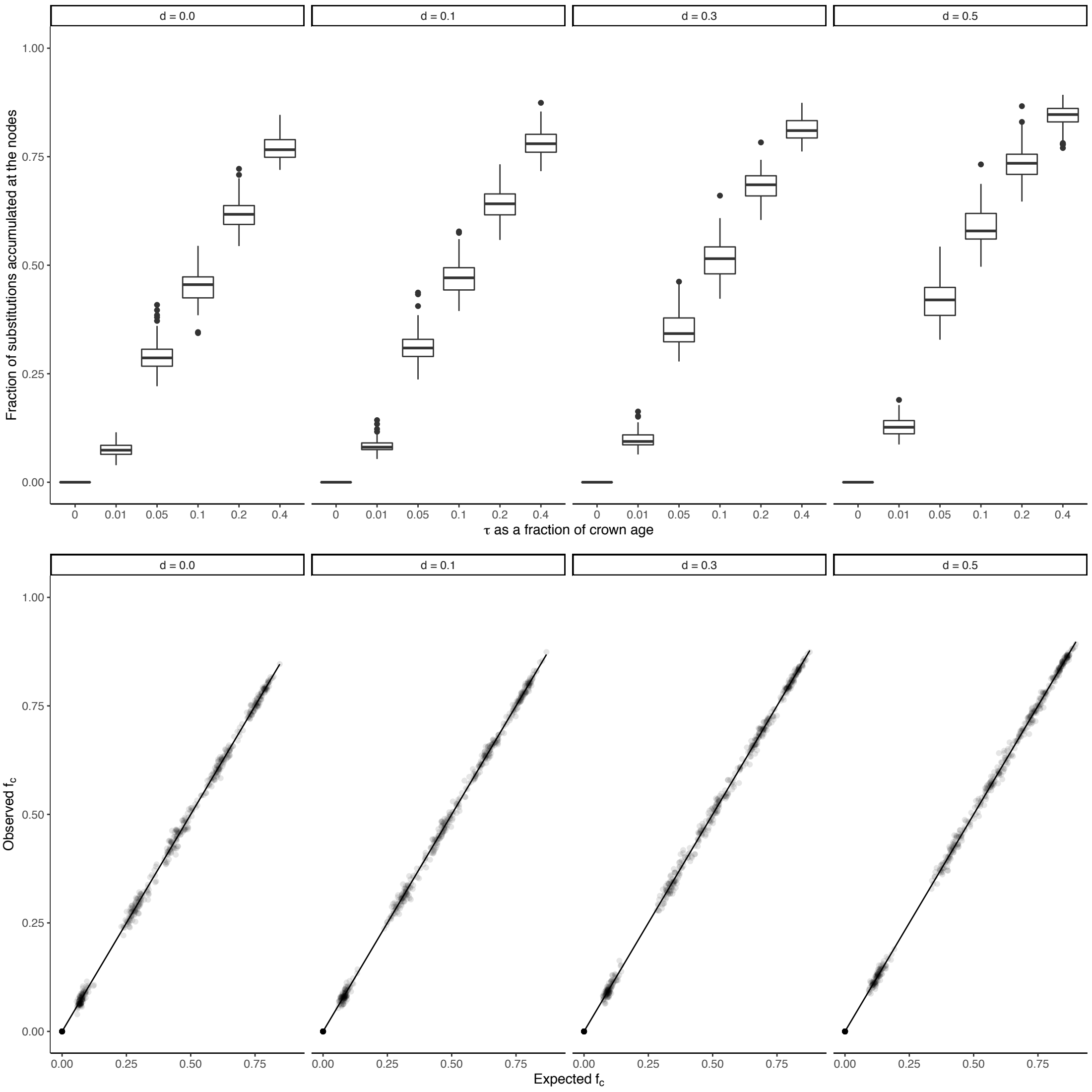


Figure S1. Top row: Observed fraction of substitutions accumulated at the nodes plotted against $\tau$ as a fraction of the crown age. Shown are results for different degrees of extinction varying in [0, 0.1, 0.3, 0.5]. Per parameter combination, 100 replicates were run. Bottom row: same parameter combinations but now showing the observed and expected $f_c$, following equation (3). The solid line indicates the observed = expected line.

Secondly, we varied the fraction $f_c$ of substitutions accumulated at the nodes compared to the total number of substitutions accumulated across the tree. We varied this fraction in [0.0, 0.1, 0.2, …, 0.9]. Using equation (3) we computed a new value for τ, given a simulated tree. For each combination of extinction rate and $f_c$ we simulated 100 true trees and generated one node substitution alignment and one *twin* alignment.

Figure S2 shows that results echo the results in the main text: with an increasing impact of node substitutions, summary statistics associated with branching times are affected most.

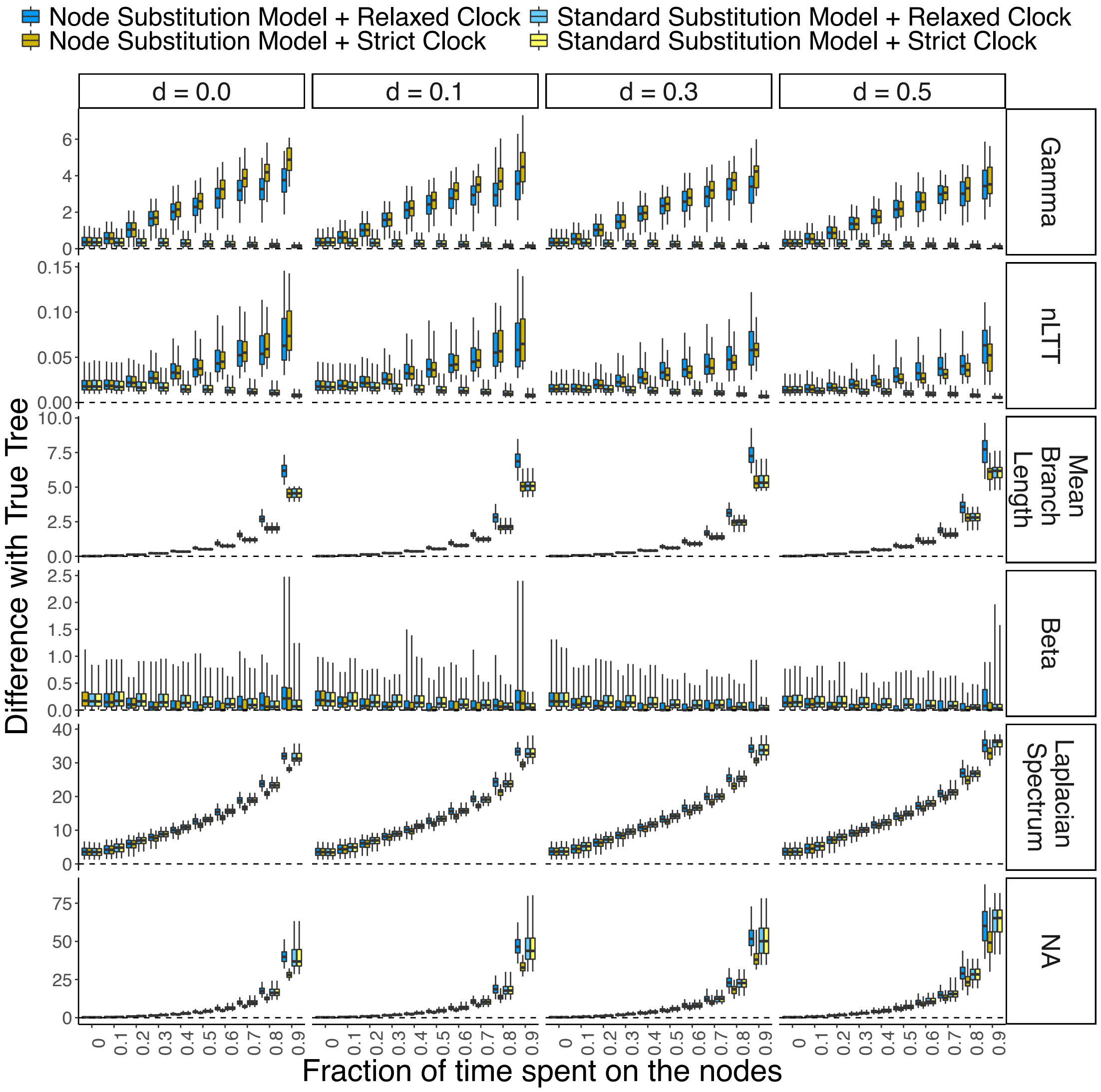

Figure S2. Difference in summary statistic values between trees inferred from an alignment generated with node substitutions (colored boxplots), and *twin* trees that were inferred from an alignment generated without node substitutions (grey boxplots). Shown is the difference with the true tree. We explore different fractions of time spent on the nodes ($\tau$, horizontal axis), and the impact of extinction ($m$, columns). The dotted line indicates zero difference between the two substitution models. Shown are results for the beta and gamma statistic, the Laplacian spectrum, the mean branch length, the nLTT statistic and crown age.

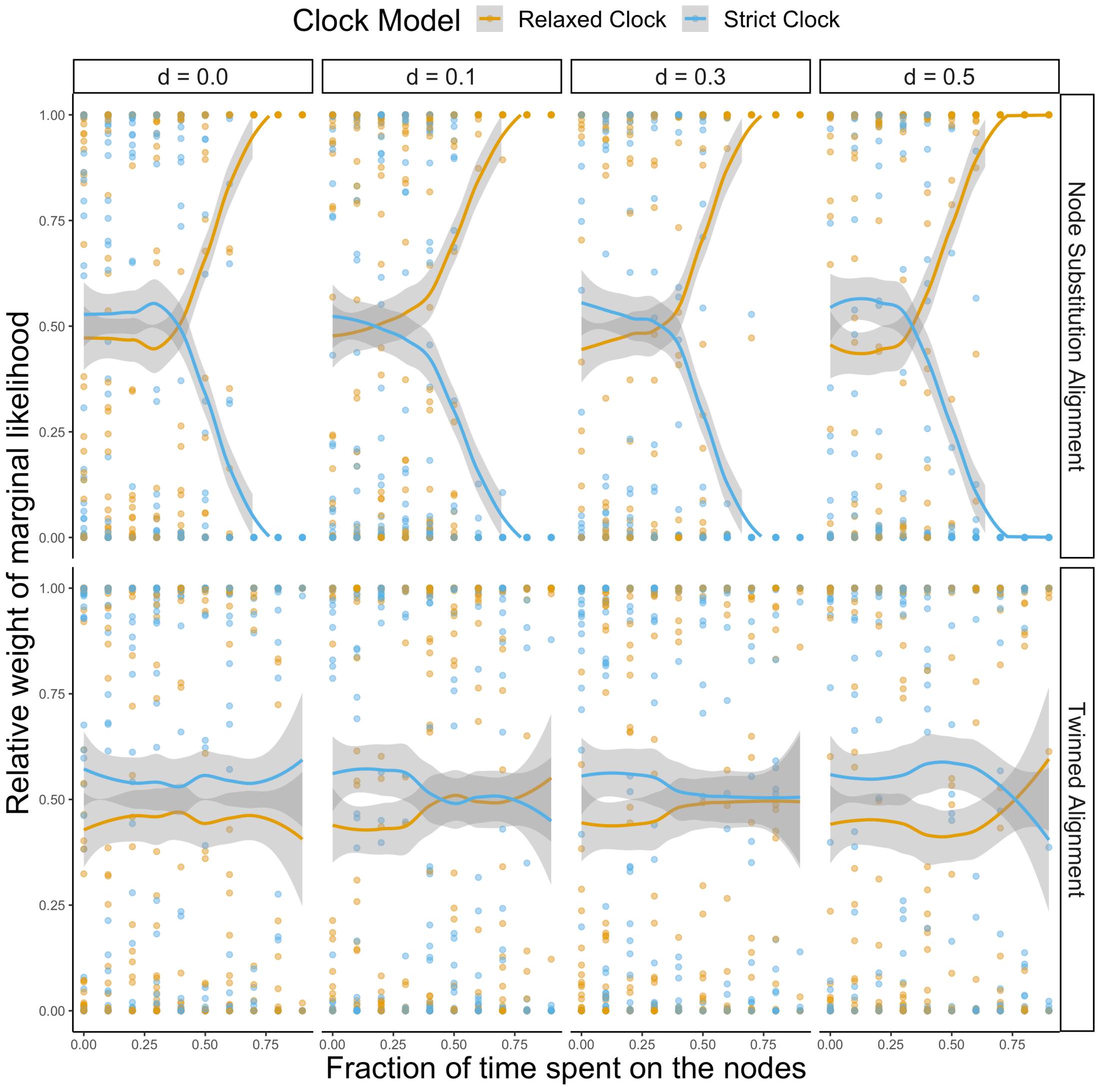


Figure S3. Marginal likelihood weight of the relaxed and strict clock model, when tested on alignments generated with a node substitution model (top row) or without node substitutions (bottom row). Per fraction of time spent on the nodes, 100 replicate trees were analyzed. Because many of the replicates share the same weights and are plotted on top of each other, solid lines indicate the best fitting generalized additive model (gam), and the 95% Confidence interval (grey shaded area) of the gam. When the fraction of time spent on the nodes increases, posterior support for the relaxed clock model increases, but only if the alignment was generated with a node substitution model.

*Linked model*

In addition to the model proposed in the main text, we studied a linked model, where both daughter branches accumulate substitutions dependent on each other, e.g. where accumulation of a substitution in one daughter branch excludes the other daughter from accumulating the same substitution (at the node). Similarly, this also precludes occurrence of a simultaneous mutation to the same base in both daughter sequences. The linked node substitution model reflects potential divergent sequence evolution, where evolution of a specific sequence in one daughter drives evolution of a divergent sequence in the other daughter.

The linked substitution model is only applied at the node and not at the subsequent branches. The transition matrix for the linked model is given by:

$\mathbf{Q}_l$ =

| | AA | TT | CC | GG | AC | AT | AG | TC | TG | CG |
|---|---|---|---|---|---|---|---|---|---|---|
| **A** | * | 0 | 0 | 0 | 1 | 1 | 1 | α | α | α |
| **T** | 0 | * | 0 | 0 | α | 1 | α | 1 | 1 | α |
| **C** | 0 | 0 | * | 0 | 1 | α | α | 1 | α | 1 |
| **G** | 0 | 0 | 0 | * | α | α | 1 | α | 1 | 1 |

where the letter on the left-hand side indicates the base present in the sequence before the node and the two letters on top of the table indicate the letters present in the two daughter sequences after the node. Thus, with rate one a single mutation occurs, for instance from A to AT, where one daughter branch inherits an A and the other daughter branch inherits a T. We do not keep track of the identity of the daughter branches, and mutations are distributed randomly over the two daughter branches. Similarly, with a rate of α, two mutations occur, and both daughter branches inherit a base that is different from the parental base, and that are different from each other. Lastly, we assume that daughter branches cannot mutate to the same base. To apply the linked $\mathbf{Q_l}$ matrix, we extend it with zeros to make it a square matrix, and then calculate the transition matrix $\mathbf{P_c}$, which now generates two sequences simultaneously.

Figure S3 shows results for the same analysis as performed for Figure 1 in the main text, but now using the linked model. Because the linked model generates fewer substitutions, effects are less strong, but as for to the unlinked model we find that for statistics related to branching times, with increasing values of $\tau$, the residual error increases.

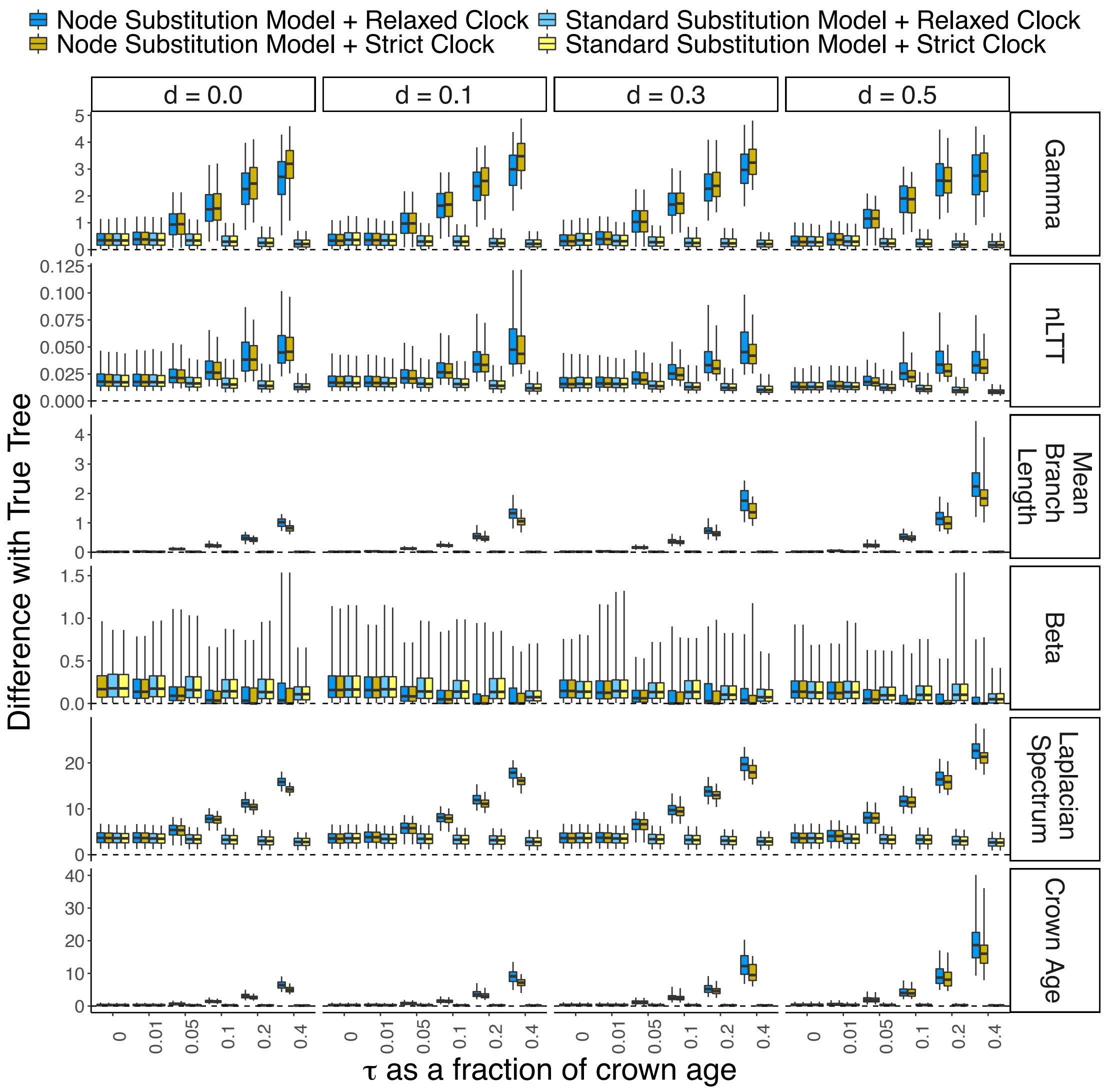


Figure S4. Difference in summary statistic values between the true tree and trees inferred from an alignment generated with node substitutions using the *linked* model (colored boxplots), and *twin* trees that were inferred from an alignment generated without node substitutions (grey boxplots). Shown are results for trees inferred using a strict clock model, and a relaxed clock model. We explore $\tau$ as a fraction of crown age (horizontal axis), and the impact of extinction ($d$, columns). The dotted line indicates zero difference with the true tree. The summary statistics are the beta and gamma statistic, Laplacian spectrum, mean branch length, nLTT statistic and crown age.

Node density effect

To assess the impact of the node density effect, we repeated the analysis from Figure 1, but using an explicit simulation framework. For this we computed the probability $P(n,i)$ of observing nucleotide $i$ at time $t$ where in total $n$ substitutions have taken place at this locus (including reverse mutations). Let us assume that we start with $i$ = A at $t = 0$, so $P_0(0,\mathrm{A}) = 1$ and $P_0(n > 0,\mathrm{A}) = P_0(n \geq 0,\neg\mathrm{A}) = 0$ (where ¬ means not, i.e. ¬A is either C, G or T). Then we can write the dynamics for $P_t(0, \mathrm{A})$ and $P_t(0, \neg\mathrm{A})$ as, dropping the subscript $t$ for notational simplicity:

$P'(0,\mathrm{A}) = -3\mu P(0,\mathrm{A})$

$P'(0,\neg\mathrm{A}) = 0$

where ʻ denotes the time derivative. The solution of these equations for the abovementioned initial condition is $P(0, \mathrm{A}) = \exp(-3\mu t)$ and $P(0, \neg\mathrm{A}) = 0$.

The equations for $P(1, \mathrm{A})$ and $P(1, \neg\mathrm{A})$ are: $P'(1, \mathrm{A}) = 3\mu P(0, \neg\mathrm{A}) - 3\mu P(1, \mathrm{A})$

$P'(1, \neg\mathrm{A}) = 2\mu P(0, \neg\mathrm{A}) + \mu P(0, \mathrm{A}) - 3\mu P(1, \neg\mathrm{A})$

We can substitute the solutions for $P(0,\mathrm{A})$ and $P(0, \neg\mathrm{A})$ here and use the initial condition to find that $P(1, \mathrm{A}) = 0$ and $P(1, \neg\mathrm{A}) = \mu t \exp(-3\mu t)$.

Similarly the equations for $P(2, \mathrm{A})$ and $P(2, \neg\mathrm{A})$ are:

$P'(2, \mathrm{A}) = 3\mu P(1, \neg\mathrm{A}) - 3\mu P(2, \mathrm{A})$

$P'(2, \neg\mathrm{A}) = 2\mu P(1, \neg\mathrm{A}) + \mu P(1, \mathrm{A}) - 3\mu P(2, \neg\mathrm{A})$

for which the solution is: $P(2, \mathrm{A}) = 3/2\ \mu^2 t^2 \exp(-3\mu t)$ and $P(2, \neg\mathrm{A}) = \mu^2 t^2 \exp(-3\mu t)$.

We can continue in this way and find that the equations are given by

$P'(n, \mathrm{A}) = 3\mu P(n - 1, \neg\mathrm{A}) - 3\mu P(n, \mathrm{A})$

$P'(n, \neg\mathrm{A}) = 2\mu P(n - 1, \neg\mathrm{A}) + \mu P(n - 1, \mathrm{A}) - 3\mu P(n, \neg\mathrm{A})$

and the general solution is:

$P(n, \text{A}) = a_n (\mu t)^n \exp(-3\mu t)$

$P(n, \neg\text{A}) = b_n (\mu t)^n \exp(-3\mu t)$

where

$a_n = 3b_{n-1} / n$

$b_n = (a_{n-1} + 2b_{n-1}) / n$

with $a_0 = 1$ and $b_0 = 0$.

We used these equations to generate sequences and then repeated the analysis as performed for Figure 1 in the main text, now knowing the exact number of substitutions that occurred. Figure S5 shows that results obtained are highly similar to the patterns found in the main text, demonstrating that the results in the main text correct adequately for the node-density-effect.

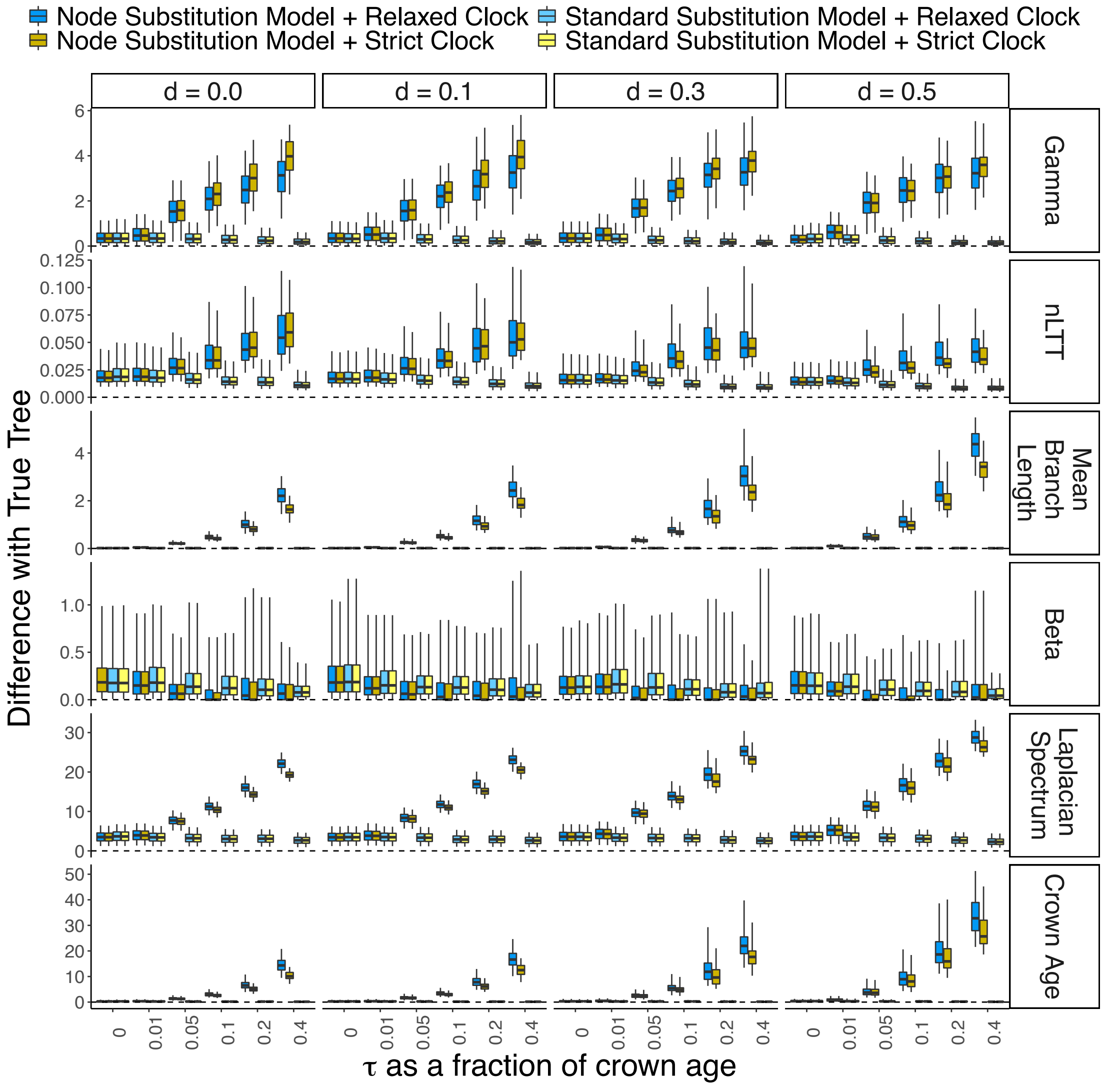


Figure S5. Identical to Figure 1 in the main text, except that mutations were simulated using the framework described above, both for sequences simulated with node substitutions and sequences simulated using the model with substitutions only on branches.
